# Interactive electrical behaviour in mormyrid weakly electric fish: Jamming avoidance response or social interaction?

**DOI:** 10.1101/2025.02.24.639824

**Authors:** Nils Weimar, Moritz Maxeiner, Emily M Wong, Ingolf P Rick, Mathis Hocke, Tim Landgraf, Gerhard von der Emde

## Abstract

Weakly electric fish emit electric organ discharges (EODs) for both active electrolocation and electrocommunication. In African mormyrids, pulse-type EODs are produced at highly variable inter-discharge intervals (IDIs), forming sequences that can convey contextual and behavioural information to conspecifics. Neighbouring fish frequently engage in time-locked interactive behaviours, such as fixed-latency echo responses and EOD synchronisation. These behaviours have been proposed to function either as a jamming avoidance response (JAR), preventing simultaneous discharges and interference with electrolocation, or as a form of social signal. To test these hypotheses, we analysed interactions between pairs of *Mormyrus rume proboscirostris*, quantifying EOD synchronisations, echo events, and instances of jamming. Our results show that jamming is rare in this species. We found no correlation between jamming frequency and echoing behaviour, nor evidence that synchronisation and echo events are direct responses to recent jams. Instead, these interactive behaviours were associated with specific movement patterns and social contexts. Synchronisations were mainly initiated by stationary hovering individuals, while echo responses were more frequent in smaller fish actively following a social partner, suggesting a role in leader-follower dynamics. Pairs with more frequent echoing also exhibited reduced inter-individual distances, indicating that interactive electrical signalling promotes social cohesion. Rather than mitigating jamming, interactive electrical behaviours in mormyrids likely serve as communicative strategies to maintain group coherence or allocate social attention. These findings highlight the role of electrocommunication in structuring social interactions. Investigating the rules of these behaviours could eventually decipher which specific IDI patterns signal social intent and help to understand the underlying mechanisms of electrocommunication.

**Highlights:**

1. Mormyrid fish frequently perform interactive signalling during electrocommunication.
2. Interactive electrical signalling is associated with specific locomotor behaviours.
3. Echo responses and signal synchronisations are not linked to jamming.
4. Signal synchronisations may support mutual assessment upon encounters.
5. Interactive signalling promotes social cohesion and leader-follower dynamics.

## Introduction

For animals living in environments where vision is limited - such as during the night or in habitats with murky waters - orientation and intraspecific communication pose significant challenges (Lissmann, 1958; Worm et al., 2021). African mormyrid weakly electric fish have adapted to these constraints by evolving an active electric sense for electrocommunication and for detecting and analysing objects in their environment, a process called active electrolocation (Lissmann & Machin, 1958; von der Emde et al., 2010). As pulse-type fish, mormyrids produce brief electric pulses at irregular intervals using a tail-based electric organ (Moller, 1995) derived from former muscle cells (Patterson & Zakon, 1997). Fish use their electric organ discharges (EODs) to navigate and detect objects such as prey items whose electrical properties differ from those of the surrounding water. In the self-generated three-dimensional electric dipole field around the fish’s body, these objects induce local modulations (Lissmann and Machin, 1958; von der Emde, 1999; von der Emde et al., 2010), which are registered by an endogenous arrangement of epidermal electroreceptor organs, the so-called mormyromasts (Bell et al., 1989; Hollmann et al., 2008).

Using a second type of electroreceptors, the knollenorgans, mormyrids can perceive and analyse foreign EODs from nearby weakly electric fish, enabling intraspecific communication (Derbin & Szabo, 1968; Baker et al., 2013). Mormyrid species that form social groups in the wild are thought to use electrocommunication to engage in coordinated behaviours in complete darkness (Arnegard & Carlson, 2005). While the EOD waveform is rather constant and allows mormyrids to exchange information on identity (i.e. about species, sex or individuality; Hopkins & Bass, 1981; Nagel et al., 2018) and relative rank in a social hierarchy (Carlson et al., 2000; Terleph & Moller, 2003), the inter-discharge intervals (IDIs) are highly variable in duration. During electrocommunication, IDIs can rapidly vary between consecutive EODs of the same individual, creating temporal IDI sequences that convey contextual and behavioural information to nearby conspecifics (Carlson & Hopkins, 2004; Baier & Kramer, 2007; Gebhardt et al., 2012a). For example, *Mormyrus rume* frequently produces so-called double pulses, i.e. alternations between short and long IDIs, as an initial threatening signal during territorial displays (Worm et al., 2017). Escalated agonistic interactions, such as physical confrontations, were found to be accompanied by discharge accelerations, often preceded by brief discharge cessations (Gebhardt et al., 2012a).

Additionally, mormyrids frequently engage in episodes of interactive electrical signalling by responding to conspecific EODs with their own discharges at a fixed preferred latency of a few milliseconds (Bell et al., 1974; Russel et al., 1974). Although these ‘echo responses’ seem to be universal among mormyrids, their behavioural significance remains unclear (Moller, 1995; Worm et al., 2018). Continuous mutual echoing between two individuals during a social encounter results in a ‘synchronisation’ of their EODs (Worm et. al, 2018), a time-locked event similar to the duetting of birds (Thorpe et al., 1972) and considered to be the most rapid form of communication in the animal kingdom (Kramer, 1990). This discrete interactive electrical behaviour may function as a communicative strategy to promote group coherence, such as when solitary individuals rejoin their group after temporary separation (Arnegard & Carlson, 2005). The ‘social attention hypothesis’ of Worm et al. (2018) proposes that a mormyrid engages in mutual echoing specifically to address another individual and allocate social attention within the electrically noisy environment of a group. This is further supported by Gebhardt et al. (2012a) who observed that synchronisation episodes can rapidly shift between different group members.

Since weakly electric fish use their EOD for both active electrolocation and electrocommunication, a conflict of interest may arise, in which social interactions between two or more fish could lead to mutual interference of their electrolocation signals (Capurro & Malta, 2004). Because of this, time-locked interactive electrical behaviours of mormyrids, such as echo responses and synchronisations, have also been hypothesised to prevent electrical jamming of the mormyromast electroreceptors during active electrolocation (Heiligenberg et al., 1978a; Moller, 1995; Capurro et al., 1999).

In the convergently evolved South American weakly electric fish species (Gymnotiformes), the jamming avoidance response (JAR) is a well-studied reflex-like electrical behaviour, in which fish shift their discharge frequency away from similar frequencies of neighbors (Watanabe & Takeda, 1963; Bullock et al., 1972; Heiligenberg et al., 1978b; Heiligenberg, 1980; Kramer, 1985; Kawasaki, 1993). Gymnotiform wave-type species such as *Eigenmannia virescens* or *Apteronotus leptorhynchus* produce continuous, quasi-sinusoidal wave-like EODs that have highly stable individual frequencies but vary across different individuals of the same species (Kramer, 1987; Shifman & Lewis, 2018). Nearby conspecific’s EODs with a similar frequency disrupt the performance during active electrolocation by causing amplitude and frequency modulations of the resulting summed signal (Heiligenberg et al., 1978b; Shifman & Lewis, 2018). To avoid jamming, individuals of wave-type species such as *Eigenmannia* rapidly shift their own EOD frequency away from the foreign signal until a frequency difference of 10 to 20 Hz has been reached (Heiligenberg 1980). The JAR helps to prevent interference of both EODs and to maintain efficient active electrolocation (Kramer, 1985), and has also been observed in the only known African wave-type species, *Gymnarchus niloticus* (Bullock et al., 1975; Heiligenberg et al., 1978b; Kawasaki, 1993).

Similarly, in pulse-type electric fish, jamming occurs when the discharges of two closely spaced individuals coincide (Capurro & Malta, 2004). While occasional interferences were found to be well tolerated, active electrolocation deteriorates quickly after multiple successive EODs have been jammed (Heiligenberg, 1974, 1976, 1980; Heiligenberg et al., 1978a). This poses a problem for South

American pulse-type fishes, which, unlike their African counterparts, emit brief EODs at intervals of low variability (Moller, 1995; Capurro & Malta, 2004). As a result, pulse sequences with similar frequencies in two interacting fish risk prolonged coinciding, potentially jamming several successive EODs. To avoid this, gymnotoid pulse species exhibit a form of jamming avoidance behaviour by temporarily shortening their IDIs to offset their EOD sequences from those of a conspecific (Heiligenberg et al., 1978a; Capurro et al., 1999). This is similar to the JAR observed in other animals with active sensory systems, such as some species of echolocating bats. These bats adjust the timing or frequency of their biosonar signals to avoid interference from the calls of nearby individuals during foraging (Jones & Conner, 2019).

With jamming avoidance strategies being widespread in both South American wave and pulse-type weakly electric fishes, as well as in the African *G. niloticus*, similar behaviours were assumed to exist in mormyrids. However, the high variability and unpredictability of mormyrid IDIs indicates that jamming is rare, raising questions about the necessity of such strategies in these fish. Despite advances in understanding mormyrid electrical interactions, data on how often such jams occur during natural interactions and how they are linked to echo and synchronisation events remain limited. Although recent developments in automatic assignment of EODs to their respective senders during experiments hold promise for generating more data on group interactions (Pedraja et al., 2021), the unpredictability of individual EODs still requires time-intensive manual analysis in current studies. This constraint has restricted previous research to a small subset of natural social interactions (Carlson, 2002; Gebhardt et al., 2012a; Worm et al., 2018).

In this study, we address this gap by analysing 35 minutes of interactions across multiple fish pairs, and explore the two possible functions of interactive electric signalling (i.e. echo responses and synchronisations) in the mormyrid, *M. rume*. To examine a potential relationship between jamming and interactive signalling, we test two distinct hypotheses: (1) If echo responses effectively mitigate jamming, there should be a negative association between jamming frequency and the time fish spent echoing, with fish pairs exhibiting more echo responses experiencing fewer jams; and (2) if echo and synchronisation events are direct avoidance responses to recent jams, the majority of these electrical interactions should follow instances of jamming. We also explore whether interactive electric signalling is associated with social behavior, investigating echo and synchronisation events in the context of the ‘social attention hypothesis’ (Worm et al. 2021). If these interactions contribute to establishing communication networks, social hierarchies, or group cohesion, we predict that (3) pairs exhibiting more echo responses would also display greater locomotor coordination, reflected for example in reduced inter-individual distances. As specific movement patterns are likely involved in electrocommunication and active electrosensing (Worm et al., 2021; Engelmann et al., 2021; Skeels et al., 2023), we analysed fish locomotor behaviour to identify potential differences between situations in which two fish synchronise their EODs versus situations in which only one fish echoes the other.

## Methods

### Animals

Captive bred *Mormyrus rume proboscirostris* were used in this study. They were kept in groups of 10 individuals in large 400-500 L holding tanks, equipped with shelters, plants and gravel. We provided a 12:12 h light:dark regime, and maintained water temperature at 25 ± 1 °C and water conductivity around 150 µS/cm. Animals were fed at least five times per week *ad libitum* with defrosted chironomid larvae. Fish were of approximately 2 years of age and measured 7.95 to 14.41 cm in standard length (SL = from the tip of the snout to the caudal base). Their sex could not be reliably determined due to their small size and the presumed absence of sexual maturation, as gonadal development is typically induced by seasonal changes in water conductivity (Moller et al., 2004; Schugardt and Kirschbaum, 2004).

16 individuals were randomly selected from two holding tanks and observed in pairs, with each fish used only once. Pairs were put together based on relative sizes. They included one larger and one smaller fish (mean size difference: 2.36 ± 1.74 cm) both to facilitate visual tracking and assess potential social hierarchy effects on interactions (Bell et al., 1974; Terleph, 2004).

### Ethical note

All experiments were carried out in accordance with the guidelines of German law and were approved by the state authority (Landesamt für Natur, Umwelt und Verbraucherschutz Nordrhein-Westfalen, LANUV NRW, reference number: 81-02.04.2020.A432). We followed the animal welfare regulations of the University of Bonn and the ‘Guidelines for the ethical treatment of nonhuman animals in behavioural research and teaching’ (ASAB, 2023). Additionally, we followed the recommendations outlined by Moritz et al. (2024) to identify relevant parameters for ensuring animal welfare in scientific fish husbandry.

The study involved only non-invasive behavioural observations. The only direct handling of the fish occurred when transferring them between their housing and experimental tanks, which were located in the same room to minimise transport stress. Mormyrids establish social hierarchies and may show intraspecific aggression, so housing tanks were enriched with ample shelters to allow individuals to avoid each other if necessary. To prevent unnecessary disruptions to social structures and minimise additional aggression, experimental pairs were always formed from individuals that were already co-housed, and fish were returned to their original tanks immediately after the experiments. Moonlight-LEDs were used to mimic the natural dim-light environment of mormyrids, ensuring conditions suitable for their activity patterns. Sudden light changes were avoided to prevent unnecessary stress. Throughout the study, fish welfare was continuously monitored using established visual indicators of stress, including skin color changes, increased gill ventilation, and alterations in swimming activity (Huntingford et al., 2006). Following the study, all fish remained housed at the Bonn Institute for Organismic Biology and continued participating in related research under the same ethical care standards.

### Experimental setup and procedures

Behavioural observations were conducted in a 120×100×20 cm tank mounted on an aluminum rack (Fig. 1A). To minimise disturbances and optimise contrast for video analysis, the lab was kept dark, with the setup illuminated by moonlight-LEDs. Light was evenly diffused through an opal glass plate with an aperture for the camera (Basler ace Classic acA2040-90um with Kowa Lens LM12HC F1.4 f12.5 mm) used to track fish movements at 25 fps. Water conditions matched those of the holding tanks of the fish (25 ± 1 °C, ∼150 µS/cm), with the water level kept at 10 cm. To prevent fish from clustering in the corners, flexible PVC plates were installed to round off the edges (Fig. 1B), promoting more uniform space use within the tank. Electrical signals were recorded using a multi-electrode array of four pairs of carbon electrodes evenly distributed along the tank walls. Each electrode featured a water-exposed carbon tip connected to a shielded coaxial cable and enclosed in a water-permeable PVC cage to prevent direct fish contact. Pilot recordings showed that such contact caused signal clipping, rendering the measured potentials unusable. All electrical signals were amplified (Brownlee Precision Model 440) and subsequently A/D-converted and recorded at a sampling rate of 50 kHz using our custom signal processing unit, the ElectroFish Interface (EFI). As this was the first application of the EFI in an experimental context, recordings were simultaneously verified using a standard data acquisition system (CED Power 1401, Cambridge Electronic Design) to ensure data accuracy and signal integrity.

**Figure 1:**
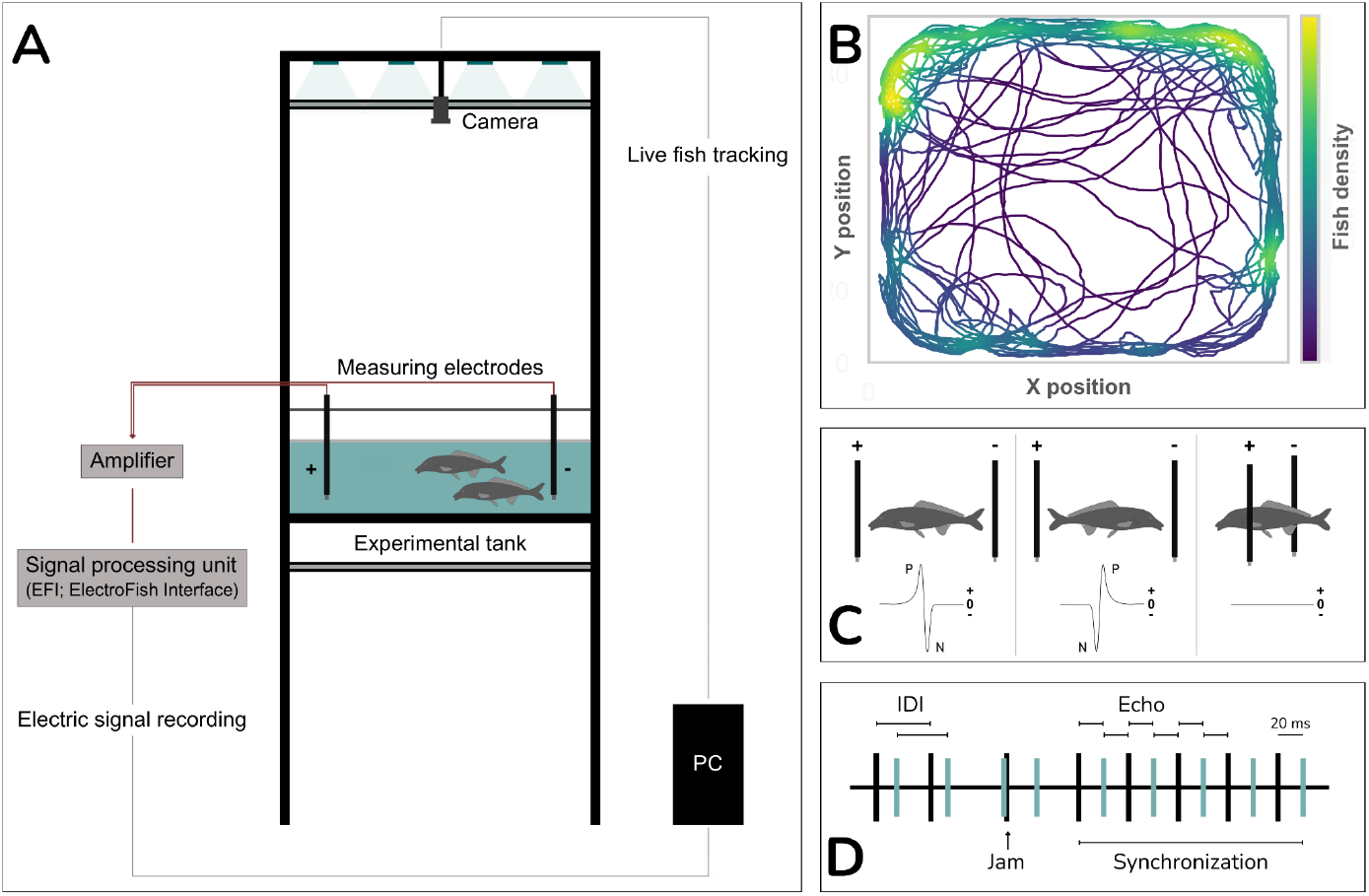
Experimental setup. **A** Tank with a multi-electrode array for recording electrical signals, amplified and subsequently processed via our custom ElectroFish Interface (EFI). The tank is illuminated with moonlight-LEDs to simulate nocturnal conditions, and swimming behaviour is tracked using an overhead camera. **B** Representative heat map showing movement patterns of a live fish pair. The colour gradient, ranging from dark blue to yellow, indicates areas of higher visitation frequency or extended duration of stay, with a preference for locations near the tank walls. Rounded gaps in the distribution outline the positions of the tank’s curved corners and the multi-electrode array, consisting of eight measuring electrodes arranged in pairs on opposite sides of the tank. **C** Appearance of recorded EOD waveforms depending on the fish’s relative position towards the electrodes. P = positive, N = negative. **D** Timeline of an exemplary electrical interaction, with vertical bars representing colour-coded EODs of two fish. The inter-discharge intervals (IDIs) denote the time between consecutive signals of the same individual. Jamming occurs when two EODs are emitted within <1.5 ms, potentially overlapping. Echo responses are characterised by a fixed latency (15-22 ms in *M. rume)* with which one fish responds to the EODs of another. Mutual echoing of both fish results in time-locked signalling sequences, described as EOD synchronisation.

Behavioural recordings were initiated by placing fish pairs inside an opaque 16 cm-diameter cylinder, randomly positioned in one of the tank corners. After a one-minute acclimation period, the cylinder was removed, and video and electrical signal recordings commenced as the fish explored the tank for five minutes. At the end of observation, fish were taken out of the tank and photographed in a transparent container on graph paper for subsequent size measurements using ImageJ. To prevent misidentification and duplicate recordings, pairs were only returned to their holding tanks after all trials were completed.

### Data analysis

Prior to analysing electrical interactions, timestamps marking the occurrence of all recorded EODs were extracted from the waveform data. Signals were then assigned to their respective senders based on amplitude and polarity relative to the fish’s positions within the multi-electrode array, as determined from the corresponding video frames (Fig. 1C). This approach, based on Gebhardt et al. (2012a), allowed for analysis of 35 minutes of video and a total of 71 867 EODs. The remaining five minutes of recordings (parts of the trials for pair #5 and pair #8) were excluded due to the fish’s similar sizes and close proximity, which made accurate EOD assignment unreliable. Trials were divided into 30-second segments for interaction analysis.

A custom Python script identified electrical jams, as well as echo and synchronisation events. Echo responses were quantified based on inter-signal intervals of EODs from different fish, previously reported in *M. rume* to occur with latencies of 15-22 ms (Carlson, 2002; Gebhardt et al., 2012a; Worm et al., 2017). To distinguish intentional echo events from accidental or undirected responses, echo events were defined as sequences of at least five echo responses (ten EODs), with no more than one non-echo interruption. Synchronisations were similarly defined as a sequence of at least five mutual echoes (Fig. 1D), also allowing a single non-echo. The durations of these electrical interactions were summed and further analysed as proportions of time within each 30-second segment, and the initiating fish (the first to echo) was identified for all events. The same script was used to detect and quantify electrical jams. Since the typical EOD of *M. rume* has a duration of ∼1.5 ms, jams were classified as two consecutive EODs from different fish with an inter-signal interval of <1.5 ms, reflecting overlap in signal duration. To assess correlations between the occurrence of electrical jams and echo or synchronisation events, a time interval of 300 ms preceding each event was examined for jams. This window, based on the average IDI of 59.51 ms calculated across all fish and recordings, typically included about five EODs preceding an event.

To synchronise recorded video and positional data with electrical signal recordings, the CED triggered an LED light marker - visible just outside the experimental arena but within the video frame - every five seconds for one second after trial initiation. Simultaneously, the CED logged timestamps of the LED activation on its electrical recording data channels, ensuring precise temporal alignment. Electrical data of the CED and EFI was synchronised by coinciding the captured EOD sequences. Fish movements were tracked using SLEAP (Social LEAP Estimates Animal Poses), an open source deep-learning based framework for multi-animal pose tracking (Pereira et al., 2022). A seven-node skeleton model, designed for *M. rume*, was used to reliably assess Cartesian coordinates and orientations (based on head and centroid nodes) for each video frame, with manual corrections applied as needed. Inter-individual distances (IIDs) as well as swimming speeds were determined per frame using the established centroid node coordinates for euclidean distance calculations. Relative spatial orientations and positions of the fish displayed before, during (at the midpoint) and after echo and synchronisation events were analysed using the custom Python script. Behavioural patterns such as leader-follower dynamics were manually assessed from the video images.

Initial observations revealed that fish pairs frequently alternated between fast-moving exploratory swimming and rather stationary hovering, suggesting distinct locomotor patterns linked to specific behaviours. To analyse these patterns systematically, we categorised fish movements during synchronisation and echo events into two modes: ‘hovering’ and ‘cruising’. Fish swimming faster than 14.41 cm/s - equivalent to the standard length of the largest individual covered within one second - were classified as ‘cruising’. Cruising behaviour involved fast, exploratory swimming throughout the experimental tank, often characterised by broader spatial movement. In contrast, ‘hovering’ was defined by slower swimming speeds and association with specific locations. This behaviour involved localised or stationary movements, such as back-and-forth swimming or parallel and antiparallel displays, and typically occurred within electrolocation distance from objects or conspecifics (Moller, 1995). The relative orientation of fish towards a base fish was analysed by categorising their alignment into two groups: parallel and antiparallel. Orientations were defined as ‘parallel’ if the angle between the fish’s orientation and the base fish was within 0°–90° to either side, indicating a similar directional alignment. In contrast, orientations were classified as ‘antiparallel’ if the angle fell within 90°–180° to either side, representing an opposing positioning relative to the base fish. These distinctions allowed us to investigate whether synchronisation and echo events occurred in different locomotor contexts.

### Statistical analysis

All data were analysed using R 4.4.2 (R Core Team, 2024). Significance tests were two tailed and performed at a significance level of α = 0.05. Generalised linear mixed models (GLMMs) implemented in the lme4 package (Bates et al., 2015) were used to assess how the frequency of mutual EOD jamming is influenced by the proportion of time fish spent echoing, as well as the number of EODs emitted by fish per 30-second time segment. Animal ID and time segment were included as random factors in both models to account for repeated measures.

GLMMs with a binomial error distribution tested whether synchronisation and echo events were more likely to follow incidents of jamming, assessing statistically significant deviations from the expected random probability of 50%. Locomotor behaviour during interactive EOD signalling was analysed by merging classifications of cruising and hovering, as well as parallel and antiparallel displays, using the ‘cbind’ function. Synchronisation and echo events were compared, with animal ID as a random factor.

A linear mixed effect model (LME) was applied to examine the relationship between event type and the average swimming speed of the initiating fish, with animal ID included as a random factor again. Two additional LMEs were used to assess factors influencing echoing behaviour and IID within fish pairs. The first model included the proportion of time spent echoing per 30-second segment as the dependent variable and relative size class (larger vs. smaller fish within pairs) as a fixed factor. The second model examined IID per 30-second segment as a function of proportional echoing duration. Both models included pair ID and time segment as random factors. IID residuals of the best explaining model did not significantly deviate from a normal distribution, as confirmed by Lilliefors test. Data on the average swimming speed of fish and echoing proportion were Box-Cox transformed (Box & Cox, 1964) to meet normality assumptions.

Model selection followed the backward elimination procedure, sequentially removing non-significant variables based on statistical relevance. The significance of fixed effects in the GLMMs and LMEs was assessed using likelihood ratio tests, comparing full and reduced models via Chi-square tests.

To compare relative positions and IIDs of initiating fish before, during and after all synchronisation and echo events, data were pooled for all individuals, tested for normality and subsequently analysed using Friedman’s two-way ANOVA with Dunn’s Test for post-hoc pairwise comparison.

## Results

Across all fish pairs, we observed 95 episodes of EOD synchronisation, with an average duration of 0.26 ± 0.12 s (max: 0.72 s), and 550 echo events, averaging 0.37 ± 0.2 s (max: 1.59 s).

### Echoing as a jamming avoidance strategy

To assess the influence of interactive electrical behaviour on jamming occurrence, we analysed the proportion of time individual fish spent echoing their partner’s EODs during a 30-second segment. EOD synchronisation events, where both fish echo each other’s EODs, were included as echo times for both individuals. Jamming incidents (overlap of the two EODs) were quantified. Jamming frequency was not significantly influenced by the proportion of time fish spent echoing (GLMM: χ2 = 2.4469, Δdf = 1, *P* = 0.1178; Fig. 2A) but showed a significant positive association with the number of EODs emitted by individual fish per 30-second segment (GLMM: χ2 = 31.763, Δdf = 1, *P* < 0.001; Fig. 2B). Although EOD synchronisation theoretically facilitates jamming avoidance, we found no evidence linking echoing time of fish to the occurrence of jams. Instead, jamming incidents were observed more frequently with higher EOD emission rates. On average, 39.1 EODs were emitted between jams, with 15.2 jams observed per 30-second segment. Despite this frequency, synchronisation events were significantly more likely to occur without a preceding jamming incident (defined as no jamming within 300 ms prior to the synchronisation) than with one (GLMM: z = -3.671, *P* < 0.001). Similarly, echo events were predominantly initiated without prior jamming (GLMM: z = -7.243, *P* < 0.001; Fig. 2C). These findings suggest that neither echoing nor synchronisations are initiated as a reaction to jamming.

**Figure 2:**
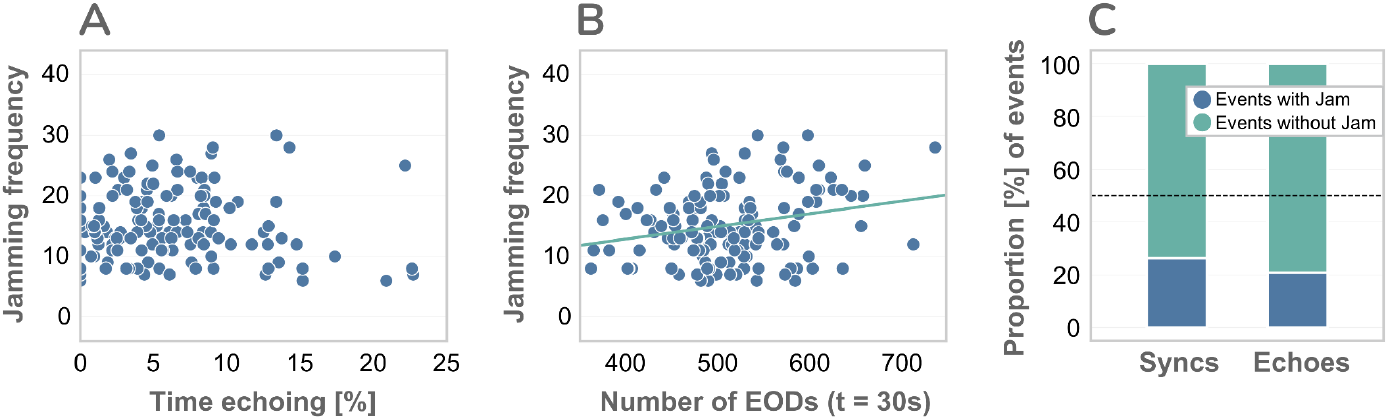
Relationship between incidents of jamming and electric behaviour in individual fish. **A** Scatter plot of the frequency of jams experienced per 30-second segment (n = 70) and the proportion (%) of time individual fish spent echoing during these segments, showing no significant correlation. **B** A significant positive association between the frequency of jams and the number of electric organ discharges (EODs) of individual fish per 30-second segment. **C** Bar graph showing the proportion (%) of synchronisation (Syncs; n = 95) and echo (Echoes; n = 550) events with (blue) or without (green) a preceding jamming event within 300 ms. A significantly higher proportion of both event types occurred without a preceding jam.

### Echoing as a social cue

Since these results show that synchronisation and echo events do not appear to function as a jamming avoidance strategy, we explored their potential role as social cues. Fish alternated between fast cruising and stationary hovering, suggesting distinct movement contexts associated with interactive electrical signalling. Locomotor behaviour differed significantly between these events (GLMM: χ2 = 10.51, Δdf = 1, *P* = 0.0012; Fig. 3). Synchronisations were predominantly associated with hovering fish, as reflected in their significantly lower swimming speeds compared to echo events (LME: χ2 = 12.723, Δdf = 1, *P* < 0.001, Fig. 4A). Echo events, in contrast, were more frequently observed when fish were actively cruising through the experimental tank (GLMM: z = -2.439, *P* = 0.0147). Notably, a significantly larger proportion of echo events were initiated by the smaller of the two involved fish (LME: χ2 = 6.3995, Δdf = 1, *P* = 0.0114; Fig. 4B), which typically occupied the follower position. As these interactions frequently occurred when one fish followed the other around the tank, EOD echoing may be indicative of distinct leader-follower dynamics and reflect social hierarchies.

**Figure 3:**
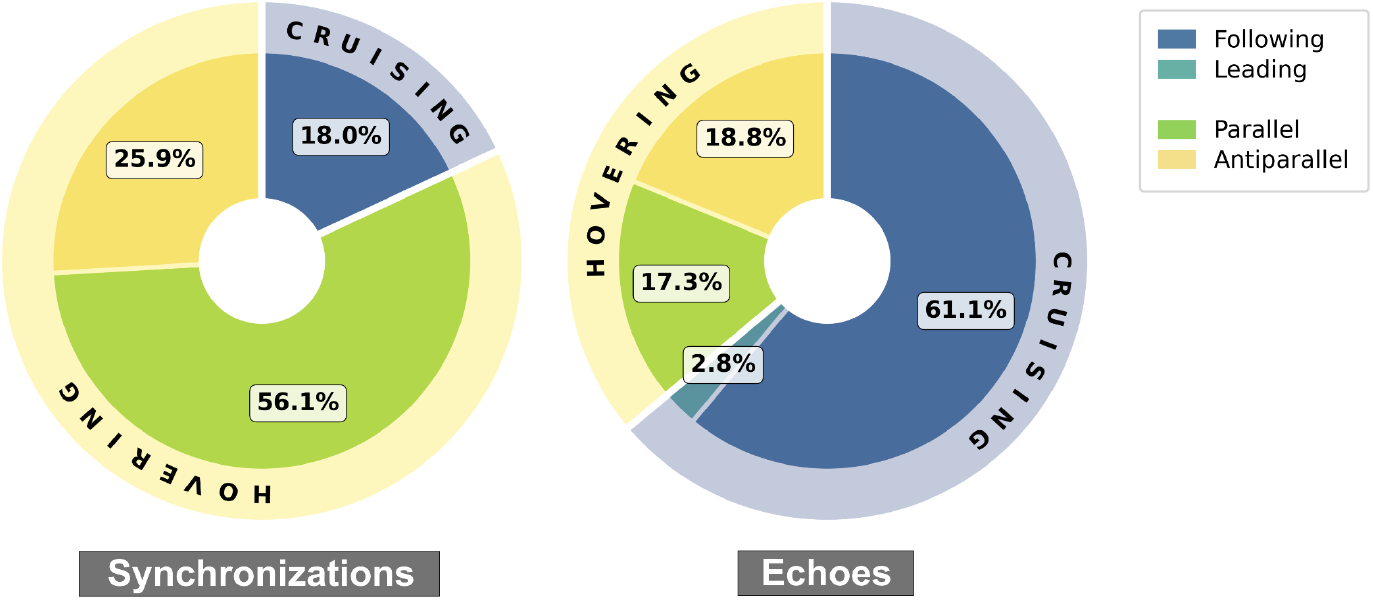
Movement patterns and interaction dynamics of weakly electric fish initiating synchronisation (left) and echo (right) events. Outer rings show the proportions of events where fish were either ‘cruising’ (light blue) or ‘hovering’ (light yellow). Inner rings indicate interaction dynamics, including whether fish were ‘following’ or ‘leading’ a social partner while ‘cruising’, and whether they were positioned ‘parallel’ or ‘antiparallel’ towards their partner while ‘hovering’. Exhibited locomotor behaviours differed significantly between event types, with synchronisation events primarily involving hovering fish and echo events associated with cruising fish.

**Figure 4:**
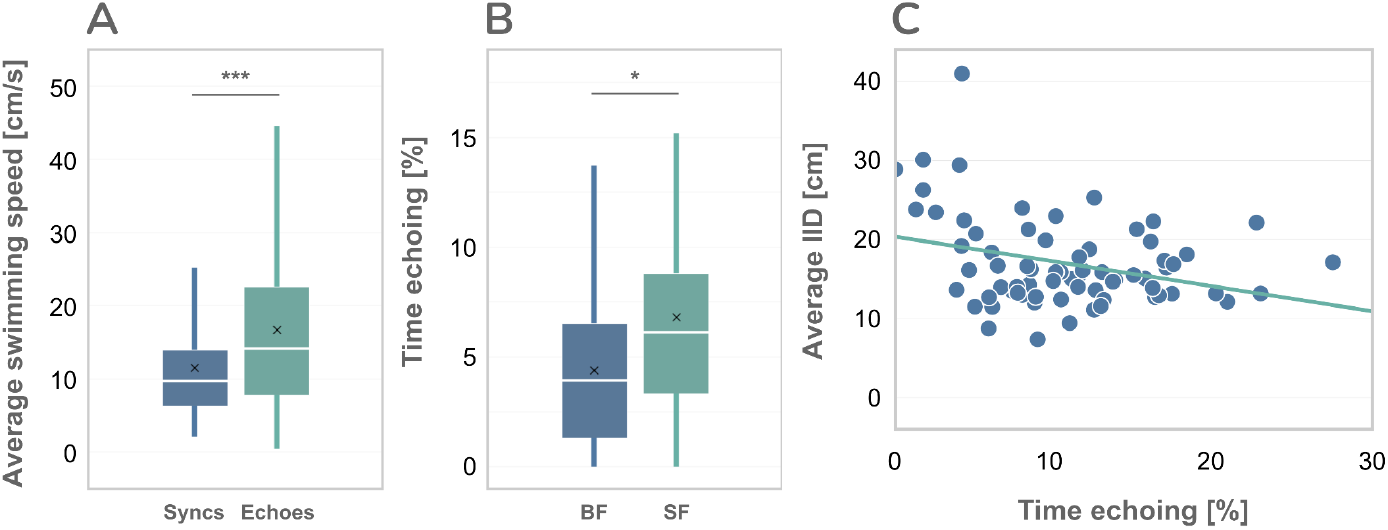
Comparative analysis of locomotor behaviours and echoing dynamics in *Mormyrus rume*. **A** Average swimming speeds (in cm/s) of the initiating fish during synchronisation (Syncs; n = 95) and echo (Echoes; n = 550) events. Non-transformed data are shown for reasons of clarity. **B** Proportions (%) of time fish spent echoing per 30-second segment (n = 70), separated by relative size class within each pair: bigger fish (BF) and smaller fish (SF). Non-transformed data are shown. **C** Scatter plot showing a significant negative correlation between the average inter-individual distance (IID, in cm) and the proportion (%) of time fish spent echoing per 30-second segment, illustrating the relationship between spatial proximity and interactive signalling. | Asterisks above the boxplots denote statistical significance: \**P* < 0.05, \*\*\**P* < 0.001.

To investigate possible spatial preferences during social interactions, we analysed the orientations of fish initiating interactive electrical behaviours. Hovering fish were categorised as being positioned either ‘parallel’ or ‘antiparallel’ based on the previously defined criteria (Fig. 3). Cruising fish were always classified as ‘parallel’ due to their leader-follower dynamics and shared orientation. We found that fish were primarily positioned parallel during interactions, with no significant difference between synchronisation and echo events (GLMM: χ2 = 2.513, Δdf = 1, *P* = 0.1129).

We also calculated IIDs between paired fish for each 30-second segment to assess their relation to echoing behaviour. Pairs that spent more time echoing exhibited significantly lower IIDs compared to those that echoed less (LME: χ2 = 8.1895, Δdf = 1, *P* = 0.0042, Fig. 4C). Further analysis revealed that fish were closest to each other during electrical interactions, with the initiating fish primarily positioned closely behind its social partner (Fig. 5). For both synchronisation and echo events, IIDs differed significantly across the Pre-Event (one second before), During Event (midpoint), and Post-Event (one second after) phases (Friedman’s two-way ANOVA; synchronisation: χ2= 28.43, *P* < 0.001; Echo: χ2 = 62.692, *P* < 0.001). Pre-Event and Post-Event IIDs were significantly higher than those during interactions (Table 1). Additionally, Post-Event IIDs were significantly lower than Pre-Event IIDs in both event types, suggesting a potential lasting effect of these interactions on social cohesion.

**Table 1:** Results of Dunn’s Test calculated for post-hoc pairwise comparison of inter-individual distances between two fish before (Pre), during, and after (Post) synchronisation and echo events.

|  | Synchronisations |  | Echoes |  |
| --- | --- | --- | --- | --- |
| | Friedman's two-way ANOVA: $\chi^2 = 28.43$, $P < 0.001$ | | Friedman's two-way ANOVA: $\chi^2 = 62.69$, $P < 0.001$ | |
| Categories | Z value | P value | Z value | P value |
| Pre - During | 4.59e-8 | < 0.001 *** | 1.97e-12 | < 0.001 *** |
| During - Post | 6.95e-4 | < 0.001 *** | 4.58e-4 | < 0.001 *** |
| Pre - Post | 1.44e-1 | 0.0481 * | 2.01e-3 | < 0.001 *** |

**Figure 5:**
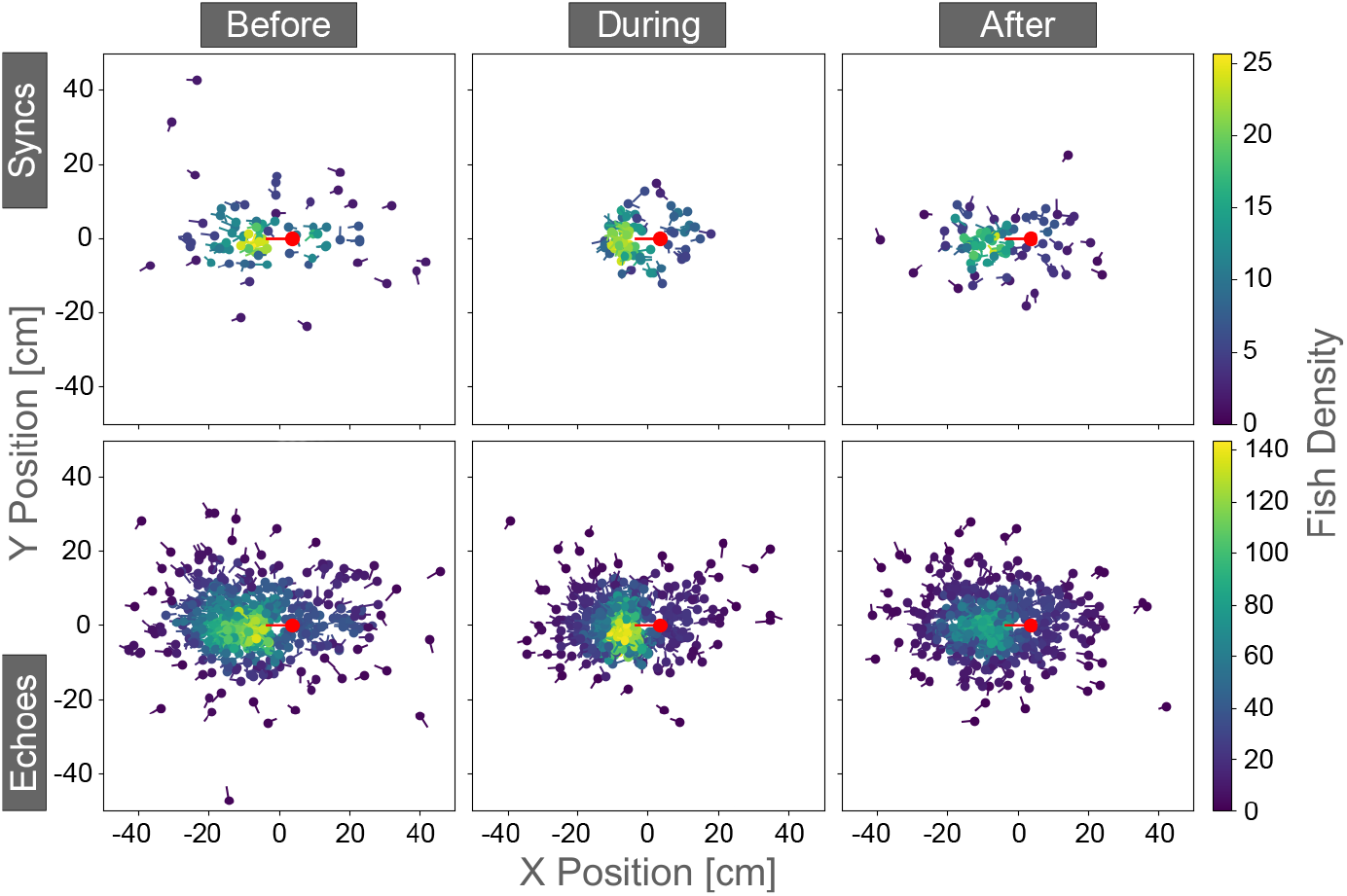
Relative XY positions (in cm) and orientations of fish initiating synchronisation events (Syncs; top row, n = 95) or echo events (Echoes; bottom row, n = 550) with respect to a partner base fish (red). Positions are depicted at three points in time: one second before (Pre-Event), during, and one second after the event (Post-Event). The density gradient illustrates spatial patterns, with lighter yellow regions indicating higher densities of relative positions. The data indicate that mean inter-individual distance is lowest during the events, with the majority of interactions initiated by fish positioned closely behind their social partner, suggesting a consistent spatial preference that may facilitate these interactions.

## Discussion

In this study, we investigated the role of interactive electrical signalling in mormyrids, focusing on electric organ discharge (EOD) synchronisations and echo events. Our results challenge the hypothesis that these behaviours primarily function as a jamming avoidance response (JAR). Instead, we propose that they serve social functions, particularly in establishing communication networks and leader-follower dynamics. This contrasts with the traditional view of jamming avoidance as an important driver of interactive electrical behaviours. Contrary to our first two hypotheses, which tested the association of jamming and interactive signalling behaviour, we found no correlation between jamming frequency and echoing behaviour, nor evidence that synchronisation and echo events are direct responses to recent jams. While the classical JAR is well-documented in South American weakly electric fish, which rely on highly regular pulse frequencies, African mormyrids exhibit more variable inter-discharge intervals (IDIs). This reduces the likelihood of EOD coincidences, which suggests that the active electrosensory system of mormyrids is less susceptible to interference, potentially eliminating the need for specialised jamming avoidance mechanisms.

Similar to our findings in *M. rume*, a study on pairs of *Brienomyrus niger* interacting in an experimental setup found that these fish experience only a few jams per 1000 pairs of EODs, suggesting that jamming is a rare occurrence for mormyrids in general (Heiligenberg, 1976). Additionally, Heiligenberg (1976) calculated that in shoals of *B. niger* with eight individuals, at most every 10th EOD of an individual was jammed. This is probably the highest level of noise these fish may naturally encounter, as only the closest neighbours produce sufficiently strong pulses to potentially interfere with electrolocation. It was proposed that mormyrids require four to eight uncontaminated successive EODs for effective electrolocation (Heiligenberg, 1976; Moller, 1995). However, Schumacher et al. (2016) demonstrated that *Gnathonemus petersii* could reliably discriminate objects even under high-frequency artificial jamming and complete EOD overlap caused by electrical white noise. None of the tested fish exhibited changes in their pulsing behaviour, indicating that *G. petersii* maintains active electrolocation even under extreme jamming. This suggests that mormyromast receptors are highly resilient to interference and that occasional natural jamming is unlikely to warrant a dedicated jamming avoidance strategy.

Interactive EOD echoing and synchronisation have been observed exclusively between two neighboring fish within a shoal at a time (Bell et al., 1974; Arnegard & Carlson, 2005; Gebhardt et al., 2012a; Pannhausen et al., 2018). In our study, an average of 39.1 EODs were emitted between jams, indicating a low probability of mutual jamming in such pairs. A jamming avoidance strategy limited to only two fish therefore seems improbable, especially given the increased likelihood of jamming with the electrical noise generated by larger groups. This brings attention to the only known example of group-level interactive electrical behaviour, described by Gebhardt et al. (2012a, 2012b), where foraging mormyrids engage in a repeating signalling pattern referred to as ‘fixed-order signalling’. In this behaviour, fish take turns generating their EODs in a set order over several rounds, potentially influenced by social hierarchies. This could mitigate jamming during high-information-value, high-EOD-rate activities like foraging (Arnegard & Carlson, 2005; Gebhardt et al., 2012a), which naturally elevate the likelihood of EOD overlap. Fixed-order signalling is characterised by highly variable inter-discharge intervals (IDIs), contrasting sharply with the time-locked nature of the mormyrid echo response. However, since we have already discussed that mormyrids likely do not require a jamming avoidance strategy to counteract mormyromast jamming and its impact on electrolocation performance, fixed-order signalling, like echoing, likely serves a different function, perhaps related to coordination or communication within a group.

Interestingly, knollenorgan electroreceptors, which are highly sensitive to conspecific EODs and are used for electrocommunication, might be more susceptible to jamming than mormyromasts. Each time a mormyrid fish produces an EOD, a corollary discharge neural signal from the EOD motor command temporarily suppresses sensory input from the knollenorgans, blocking the detection of foreign EODs during this time, possibly to prioritise mormyromast-mediated active electrolocation (Bell & Grant, 1989; von der Emde & Bell, 2002). As a result, mormyrids are temporarily insensitive to foreign pulses both during and immediately following the emission of their own EOD. This way, the dual use of the EOD for both electrolocation and electrocommunication may present a unique challenge for mormyrids, particularly in socially relevant contexts such as encounters with unfamiliar conspecifics. While the classical JAR in South American weakly electric fish is primarily geared towards preserving the integrity of electrolocation (Bullock et al., 1972; Heiligenberg, 1980; Moller 1995), the potential susceptibility of mormyrid knollenorgan electroreceptors - and thus electrocommunication - to jamming has received less attention. It could be that mormyrids may need a strategy to mitigate interference in scenarios requiring electrocommunication and use echoing and synchronisation behaviours to do so.

Episodes of synchronisation may represent an alternative knollenorgan jamming avoidance mechanism - hereafter termed ‘Communicative jamming avoidance strategy’ (C-JAS) - tailored for socially significant interactions, ensuring effective communication without compromising electrolocation. Echoing fish encountering one another may thus ensure that their EOD follows the reafferent 2-millisecond inhibition period of the receiver’s knollenorgans (Russel & Bell, 1978; Bell & Grant, 1989), while simultaneously minimising the rare likelihood of overlap with the receiver’s subsequent EOD even more. As a result, the echoing fish provides its social partner an opportunity to analyse its EODs, while in turn gaining the chance to assess the EODs of that partner. This aligns with observations that synchronisation events often occur when solitary fish rejoin their group after temporary separation (Arnegard & Carlson, 2005) or when fish approach a conspecific from behind (Worm et al., 2018), as also found in this study.

Our third hypothesis predicted that pairs with more frequent echo responses would exhibit greater locomotor coordination, as reflected in reduced inter-individual distances. This was indeed supported by our findings, as pairs with higher echoing frequencies demonstrated closer proximity and synchronised movement patterns. Such behaviour supports Worm et al.’s (2018) social attention hypothesis, suggesting mutual echo responses resulting in EOD synchronisation enable fish to recognise and address specific individuals within a group in order to establish brief dyadic communication frameworks. Interaction with familiar conspecifics enhances group cohesion in mormyrids, facilitating better coordination (Serrier & Moller, 1981). This way, synchronisations may function as a form of mutual greeting and assessment, serving as a precursor to the exchange of contextual information and eventually the emergence of social dynamics and coordinated group behaviours. Additionally, synchronisations may signal a willingness to cooperate and the absence of aggression, fostering non-threatening interactions. Our analysis of movement patterns during synchronisation events supports this interpretation. Synchronisations were typically initiated by fish positioned closely behind their social partner while both were hovering in place (Fig. 3 & 5). Afterwards, inter-individual distances were reduced compared to before. This suggests that the fish approach a conspecific from behind, signal non-aggressive intent through synchronisation, establish a communication channel, and then move together while maintaining close proximity. Interestingly, synchronisation events in this study were relatively rare and shorter in duration (mean: 0.26s; max: 0.72s) compared to previous reports (up to several seconds; Worm et al., 2018). This likely reflects our strict threshold for synchronisation detection, but also the experimental setup, in which each fish had only a single, constant social partner within a restricted area. As a result, the need to allocate social attention was minimal, as there were no opportunities for repeated or genuinely novel encounters.

In contrast to synchronisations, we observed substantially more episodes of unidirectional echoing. When the mormyrid echo response was first described, it was linked to ‘antiparallel’ displays during agonistic interactions (Bell et al., 1974; Moller, 1995). However, our findings reveal that only a small fraction of echo events occurred with the initiating fish positioned inversely to its social partner. Significantly more of these events took place with both fish aligned in orientation (Fig. 3), suggesting spatial preferences that may be linked to the alignment of sensory fields, as previously noted by Terleph (2004). Moreover, we observed that smaller fish spent significantly more time echoing the EODs of larger fish than vice versa. Unlike synchronisations, which predominantly occurred while both fish remained hovering, unidirectional echoing was primarily observed as the initiating fish engaged in the characteristic single-file swimming of mormyrids (Moller, 1976; Gebhardt et al., 2012a), closely following its social partner around the tank in an exploratory context. Previous studies have shown that shoaling mormyrids establish size-dependent social hierarchies (Bell et al., 1974; Terleph, 2004). Our findings suggest that echoing behaviour reflects these hierarchies, supporting Schumacher et al.’s (2016) hypothesis that echoing conspecific EODs functions as a submissive, rather than agonistic, signal.

Our results indicate that interactive electrical signalling in mormyrid weakly electric fish is not linked to the classical JAR, which is likely not required to maintain effective electrolocation. Instead, the primary function of interactive signalling may be social, potentially acting as a C-JAS and/or as an independent cue to initiate group dynamics. Mormyrids have been shown to engage in highly coordinated group behaviours, some are e.g. more successful hunting in groups (Arnegard & Carlson, 2005), and some may even be able to utilise the EODs of conspecifics to extend their electrolocation range (collective sensing; Pedraja & Sawtell, 2024). These findings highlight the importance of mechanisms that facilitate group cohesion. Notably, single-file swimming, as well as parallel orientated lineup behaviours, were previously shown to be absent in groups of operated, electrically silent mormyrids (Moller, 1976). We propose that interactive electrical signalling, such as the mormyrid echo response, may thus serve to establish and maintain communication channels, facilitate leader-follower relationships, and promote group cohesion in their dark and turbid natural environments. As pairs that spent more time echoing also maintained closer inter-individual distances, interactive electrical signaling may have lasting effects on movement patterns and spatial positioning within pairs. It would be particularly interesting to analyse the IDI sequences that follow episodes of synchronisation to explore potential patterns correlated with locomotor behaviours, such as leader-follower dynamics. For instance, specific IDI sequences might serve as indicators of which individual assumes the leading role or signal a mutual agreement to synchronise movements. These patterns could provide deeper insight into how synchronisation events establish or reinforce social roles within a pair or a group. Moreover, recurring IDI motifs could reveal how fish negotiate spatial positioning and coordination in dynamic environments. Future studies examining these sequences in greater detail, especially in more complex social settings, could illuminate the nuanced mechanisms through which mormyrids integrate electrocommunication and movement to maintain group cohesion.

## Acknowledgments

We thank Frank Kirschbaum for providing the mormyrid weakly electric fish, and Barbara Bauch and Slawa Braun for their expert care and management of the animals. This study was funded by the Deutsche Forschungsgemeinschaft (DFG, German Research Foundation) - Project numbers: EM 43/21-1 and LA 3534/4-1.

## Author’s contributions

**Nils Weimar:** Conceptualization, Methodology, Software, Formal Analysis, Investigation, Data Curation, Writing - Original Draft, Visualization. **Moritz Maxeiner:** Methodology, Software, Data Curation. **Emily M Wong:** Software, Formal Analysis. **Ingolf P Rick:** Formal Analysis. **Mathis Hocke:** Data Curation. **Tim Landgraf:** Software, Writing - Review & Editing, Supervision, Project Administration, Funding Acquisition. **Gerhard von der Emde:** Conceptualization, Writing - Review & Editing, Supervision, Project Administration, Funding Acquisition.

## Data availability

All raw video, tracking, and electrical signal data underlying the results will be publicly available in a Zenodo repository upon acceptance.

